# Live imaging the Foreign Body Response reveals how dampening inflammation reduces fibrosis

**DOI:** 10.1101/498444

**Authors:** David B. Gurevich, Kathryn E. French, John D. Collin, Stephen J. Cross, Paul Martin

## Abstract

Implanting biomaterials such as surgical sutures leads to wound inflammation and a Foreign Body Response (FBR), which can result in scarring and ultimately biomaterial rejection. To investigate the cell and signalling events that underlie FBR, we use live imaging of zebrafish reporter lines to observe how inflammation and angiogenesis differ between a healthy acute wound versus suture implantation. We observe inflammation extending from the suture margins and correlates with subsequent avascular and fibrotic encapsulation zones: sutures that induce more inflammation result in increased zones of avascularity and fibrosis. Moreover, we capture macrophages as they fuse to become multinucleate foreign body giant cells (FBGCs) adjacent to the most pro-inflammatory sutures. Both genetic and pharmacological dampening of the inflammatory response minimises the FBR (including FBGC generation) and normalises the status of the tissue surrounding these sutures. This new model of FBR in adult zebrafish allows us, for the first time, to live image the process and to modulate it in ways that may lead us towards new strategies to ameliorate and circumvent FBR in humans.

## Introduction

The surgical implantation of medical devices and biomaterials - from cardiac pacemakers to prosthetic joints to surgical sutures - has increased greatly over recent years as technologies advance and the population ages (Major et al, 2015). Once a material is implanted in host tissue, the interactions between the biomaterial and surrounding cells and matrix are critical in determining whether successful integration occurs. In ideal circumstances, biomaterial implantation results in an acute inflammatory response that drives a significant and necessary wound angiogenic response and subsequently limited fibrosis/scarring; this scenario largely recapitulates acute wound healing and leads to resolution of the repair response and successful biomaterial integration. Failure of biomaterial integration can be due to the exacerbation of the Foreign Body Response (FBR), where acute inflammation transitions to chronic inflammation and is generally accompanied by foreign body giant cell (FBGC) formation and results in fibrous encapsulation (Anderson et al, 2008). This response can limit the efficacy of implantable biomaterials, leading to rejection and adverse outcomes that impact patient quality of life and cause a significant burden on healthcare economics. Most previous studies of FBR have been performed on mammalian models such as dogs and mice, largely using histology on fixed samples as an endpoint (Klopfleisch, 2016; Selvig et al, 1998), although more recent intravital studies of implanted plastic chambers in a mouse skin fold model have enabled a degree of dynamic imaging of collagen deposition during FBR using second harmonics (Dondossola et al, 2016). However, these studies are not optimal for high resolution investigations of the multifaceted and dynamic molecular conversations that occur between tissue and biomaterial; nor can they explain how some materials integrate well with minimal scarring while others undergo an extensive FBR and are ultimately rejected.

We have developed a genetically tractable and translucent model of the FBR that allows for transgenic fluorescent marking of various cells and tissues, enabling the real-time visualisation of immune cell-foreign body interaction over time in a non-invasive manner (Witherel et al, 2017). Aside from external fibrin clot formation, most steps of mammalian wound repair appear to be well conserved in zebrafish and have previously been extensively characterised (Gurevich et al, 2018; Mathias et al, 2009; Renshaw et al, 2006; Richardson et al, 2013); all the initial tissue interactions that are believed to contribute to the development of FBR are known to be present. By fluorescently labelling leukocytes, inflammatory markers and blood vessels in the living organism, we are able to study the dynamic activities of these lineages in response to the implanted biomaterials, observing the interactions between these cells and the subsequent fibrotic encapsulation, and how these interactions can be modulated to reduce fibrosis and improve integration of biomaterials.

## Results

### The extent of non-resolving scar surrounding the foreign material varies according to suture type

Fibrotic encapsulation of foreign bodies, including biomaterials, is a key component of FBR, and is critically important in determining how well a material is integrated into the surrounding tissue (Mikos et al, 1998; Ward, 2008). Previous investigations have shown that materials vary in their biocompatibility, with a consequent variation in degree of inflammatory response and the extent to which fibrosis is induced following implantation (Bryers et al, 2012). To investigate whether zebrafish tissue exhibits a similar, variable fibrotic response during FBR to that seen in mammalian tissues, we implanted either 8-0 non-resorbable monofilament nylon or resorbable braided polyglycolic acid (vicryl) sutures of the same dimensions into flank tissue anterior to the base of the tail fin (Figure 1A, B). Control acute wounds were generated by ‘pulling through’ a vicryl suture at the same location (Figure 1B). It is already established that, unlike mammalian skin, acute wounds in adult zebrafish skin initially deposit scar collagen but this subsequently resolves (Richardson et al, 2013). Our Masson’s trichrome histological staining indicates persistent scarring and fibrosis in FBR instances by our endpoint of 28 days post suture implantation (DPS), contrasting with the resolving scarring observed in acute wound repair (Figure 1C, D, Sup. Figure 1A, and see (Richardson et al, 2013)). Importantly, the extent of the fibrotic area surrounding vicryl sutures was much larger than the response to nylon sutures (0.3297mm^2^ compared with 0.0392mm^2^, Figure 1D), suggesting that zebrafish tissues react to these materials in similar ways to mammalian tissues.

**Figure 1:**
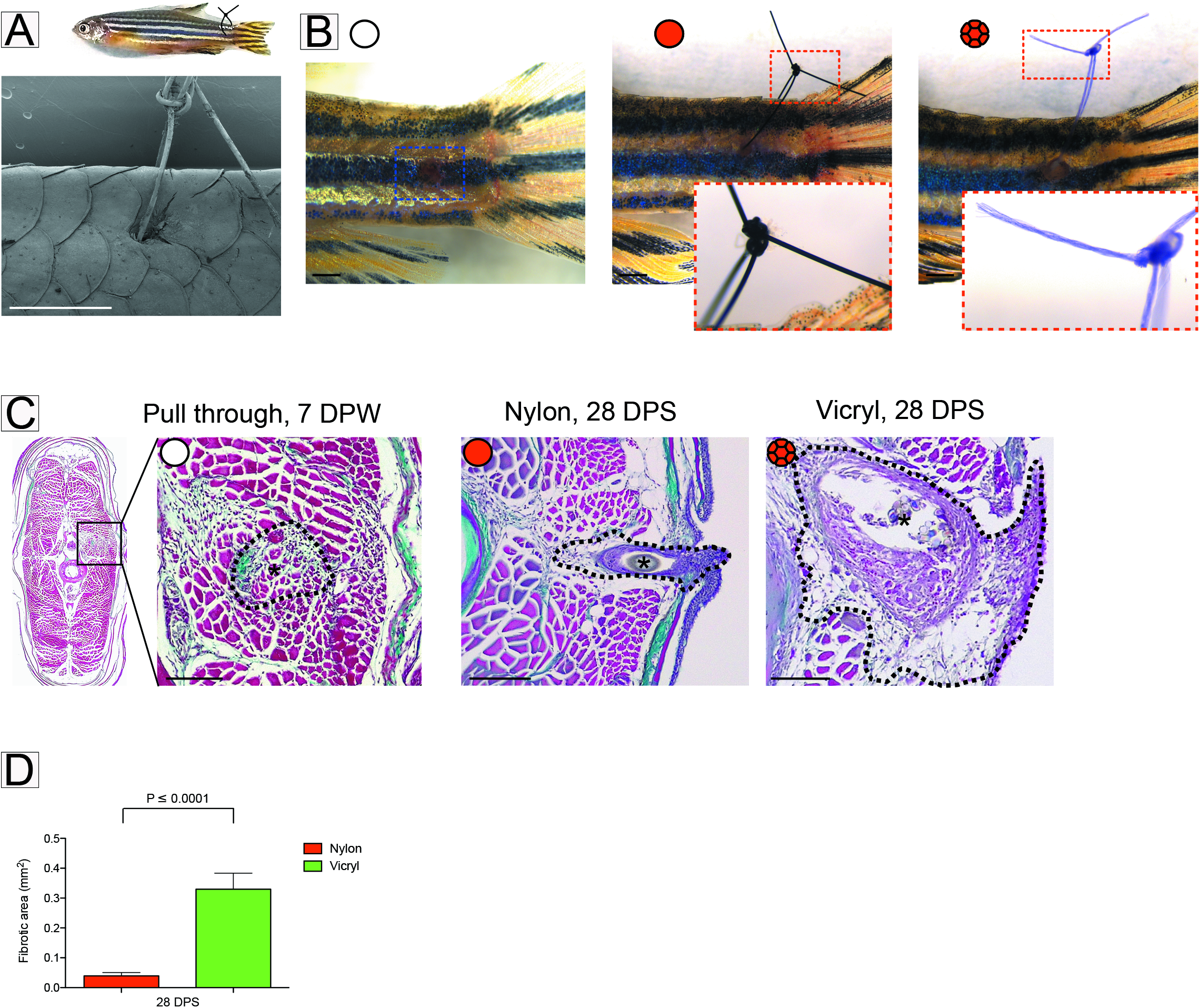
Extent of foreign body fibrotic encapsulation is dependent on suture type. A) Schematic illustration of the zebrafish suture model, with scanning electron micrograph showing a suture in place. N = 6 independent fish. B) Representative images of zebrafish following suture pull through (white circle), nylon suture (red circle) or vicryl suture (red braided circle), 1 day post suturing (DPS) with insets to show suture detail. N = 5 independent fish per condition. C) Masson’s Trichrome stained transverse sections of pull through at 7 days post wounding (DPW), nylon or vicryl sutured fish at 28 DPS, to indicate extent of fibrosis. Sutures indicated with black asterisks; zone of scarring and fibrotic encapsulation indicated by black dotted line overlay. N = 5 independent fish per condition. D) Quantification of total area of fibrotic encapsulation, measured from images in C). Statistical significance is indicated, as determined by two-tailed *t* test. Scale bars: A = 1mm, B = 1mm, C = 100μm.

### All suture types drive an exaggerated and prolonged immune cell response, and the degree of scarring correlates with the extent of inflammatory response

We next utilized the translucency of the fish to visualise the dynamic interactions underlying the establishment of the FBR. We imaged the same fish and followed the same FBR over extended time periods without the interference of a viewing chamber or other implanted intrusions. Following suture implantation, high resolution live imaging was used to view Tg(*mpx*:GFP); Tg(*mpeg*:mcherry) double transgenic zebrafish, which mark neutrophils and macrophages, respectively (Ellett et al, 2011; Renshaw et al, 2006). By imaging the same fish at specific time points across the observed 28 DPW or DPS, we were able to determine the differences in immune cell response to the two suture materials versus acute (pull through sutures) wounding, over time (Figure 2A). Previous studies examining FBR showed that the first immune cells to encounter the biomaterial are neutrophils (Selders et al, 2017), an observation supported by our results. In acute wounds, neutrophil and macrophage numbers peak at 4 DPW and 14 DPW, respectively, after which they resolve back to uninjured levels (Figure 2B). We see a similar pattern for nylon sutures, although some persistent immune cells remain in the vicinity of the suture at 28 DPS (Figure 2B). By contrast, we observe a large and persistent immune response up to 28 DPS in vicryl sutured fish, with many immune cells, particularly macrophages, remaining in close proximity to the suture edge (Figure 2A, B); this immune cell retention appears to correlate with the extensive fibrosis seen in response to this suture type (Figure 1C, D).

**Figure 2:**
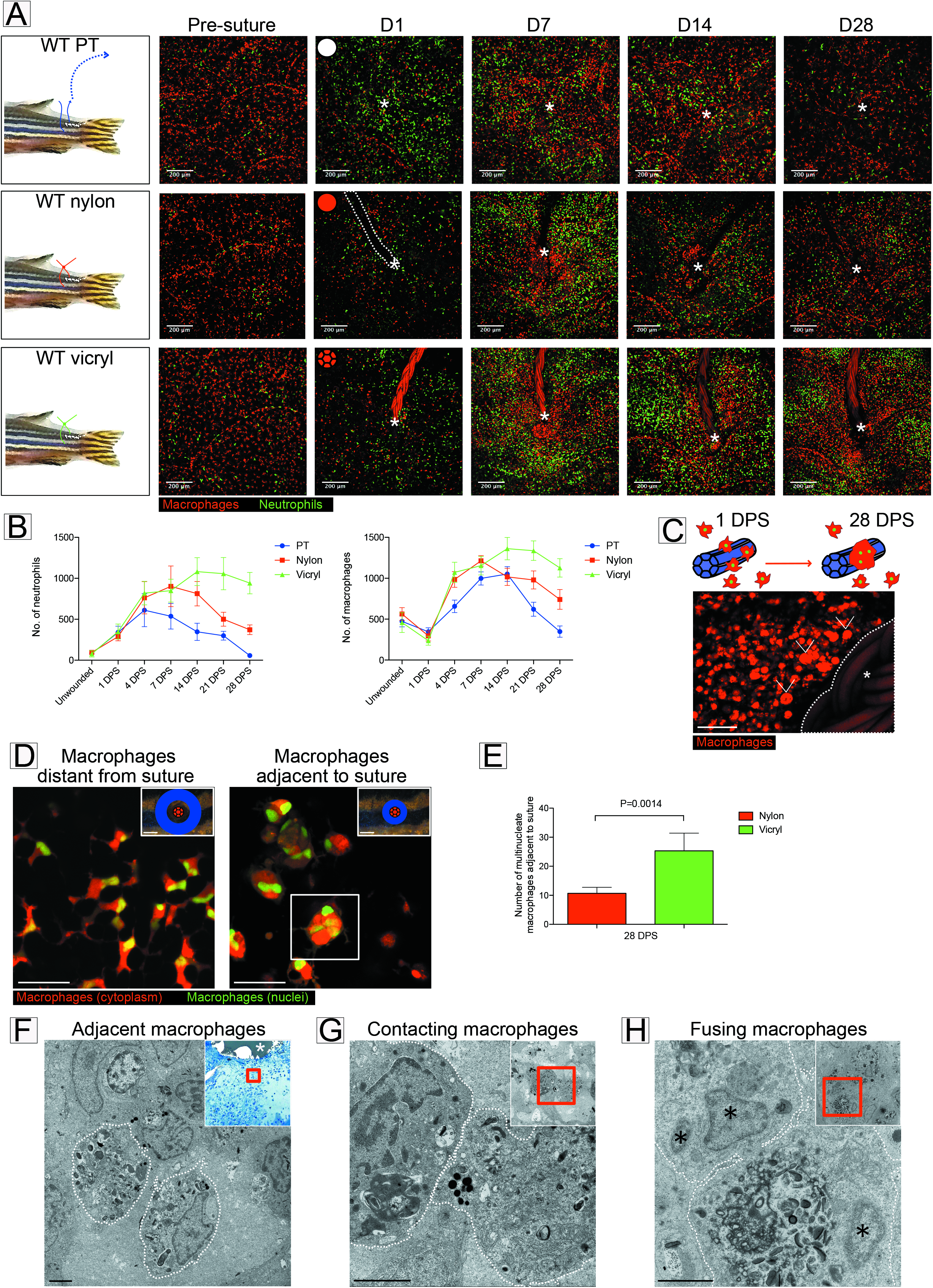
Magnitude of immune response and numbers of foreign body giant cells are greater for vicryl versus nylon sutures. A) Schematic and representative images of Tg(*mpx*:GFP); Tg(*mpeg*:mCherry) double transgenic zebrafish immediately prior to and following suture pull through, or implantation of nylon or vicryl sutures, at indicated timepoints. Area of wounding or implantation marked by white asterisk; dotted lines indicate nylon suture. N = 6 independent fish per condition. B) Quantification of neutrophil and macrophage numbers in the vicinity of wound/suture, measured from images in A). C) Schematic of macrophage fusion observed in the vicinity of the suture, and representative image of Tg(*mpeg*:mCherry) transgenic adult zebrafish at 28 DPS, showing larger “fused” macrophages, or FBGCs (arrowheads), adjacent to vicryl suture (white asterisk) with more standard-size macrophages further from the suture. N = 5 independent fish per condition. D) Representative images of Tg(*mpeg*:mCherry); Tg(*mpeg*:nlsClover) double transgenic zebrafish at 28 DPS, showing that the larger macrophages adjacent to sutures (within 200μm radius) are multinucleated (an exemplar such cell indicated by boxed area, is revisited in Sup Fig 2), compared to normal sized, single-nucleated macrophages at more distal sites (adjacent and “distal to suture” zones illustrated in blue). N = 6 independent fish per condition. E) Quantification of images from D, indicating the number of large, multinucleated macrophages adjacent to a wound/suture for nylon and vicryl sutures, at 28 DPS. Statistical significance is indicated, as determined by two-tailed *t* test. F-H) Representative electron micrographs showing three different stages of macrophage-macrophage interactions at 28 DPS: (F) nearby but not directly contacting macrophages (membranes indicated by dotted lines); (G) two adjacent macrophages come into direct plasma membrane contact; (H) a membrane fusion of two macrophages. Insets are low magnification images with red box highlighting the high magnification view. In (F) low magnification inset is a Methylene Blue stained thick section with the suture indicated by a white asterisk. In (H) macrophage nuclei are indicated by black asterisks,). N = 4 independent fish. Data information: error bars indicate mean ± SD. Scale bars: A = 200μm, C = 50μm, D = 20μm (inset = 200μm), F-H = 2μm.

Close observation of suture-associated macrophages at timepoints beyond 14 days post implantation indicates that some of these cells appear to be considerably larger than standard macrophages (Figure 2C). This is suggestive of a phenomenon seen in mammalian FBR and TB granulomas, where macrophages converge and fuse, transforming into foreign body giant cells as a consequence of chronic inflammation (Davis et al, 2002; Sheikh et al, 2015; ten Harkel et al, 2016). To examine whether a similar response to foreign body induced chronic inflammation may be occurring here, we used a Tg(*mpeg*:mCherry); Tg(*mpeg*:nlsClover) double transgenic fish to enable visualization of both cytoplasm and nuclei of macrophages (Figure 2D). By contrast to pull through injuries, where FBGCs were not detected, we observed a significant number of large, multinucleated macrophages – some with up to 6 nuclei – in close proximity (within 200μm) of the suture at 28 DPS, with larger numbers of these FBGCs around vicryl sutures than adjacent to nylon sutures (Figure 2D, E, Sup. Figure 2A, Movie 1). To complement our live imaging analyses we undertook transmission electron microscopy in an attempt to capture the moments when adjacent mononucleate macrophages fuse to generate multinucleated foreign body giant cells (FBGCs) (Figure 2F-H). Together, these results indicate that zebrafish immune cells interact with foreign bodies in very similar ways to those observed in mammalian models, and that these interactions may be a key component in directing the extent of fibrotic encapsulation in response to implanted biomaterials.

### Extent of fibrosis reflects immune cell dynamics within the suture-adjacent tissue

Having observed the association between extent of inflammation and degree of subsequent scaring at the suture site, we wondered whether live imaging of leukocyte behavior in response to the suture might provide insight into the development of the FBR. Performing timelapse photomicroscopy on Tg(*mpeg*:mcherry) unwounded zebrafish flank tissue revealed that macrophages are relatively sparse and static (Movie 2). Following acute wounding, neutrophils and macrophages rapidly migrate towards the wound site; by one day post wounding their motility is still rapid, but their movement once at the wound site lacks directionality (Figure 3A-C, Movie 3). Suture implantation led to an increase in immune cell directionality at early timepoints, but a marked reduction in speed at later timepoints, such that macrophages appear ‘paralysed’ in the tissue adjacent to the suture, particularly in the case of vicryl sutures (Figure 3B, C, Movies 4-7). This suppression of cell movement extended further from the vicryl suture than for nylon sutures and correlates with the increased extent of fibrosis associated with vicryl sutures (Figure 1D). These results suggest a causal association between inflammation and localized fibrosis that we might test in our model.

**Figure 3:**
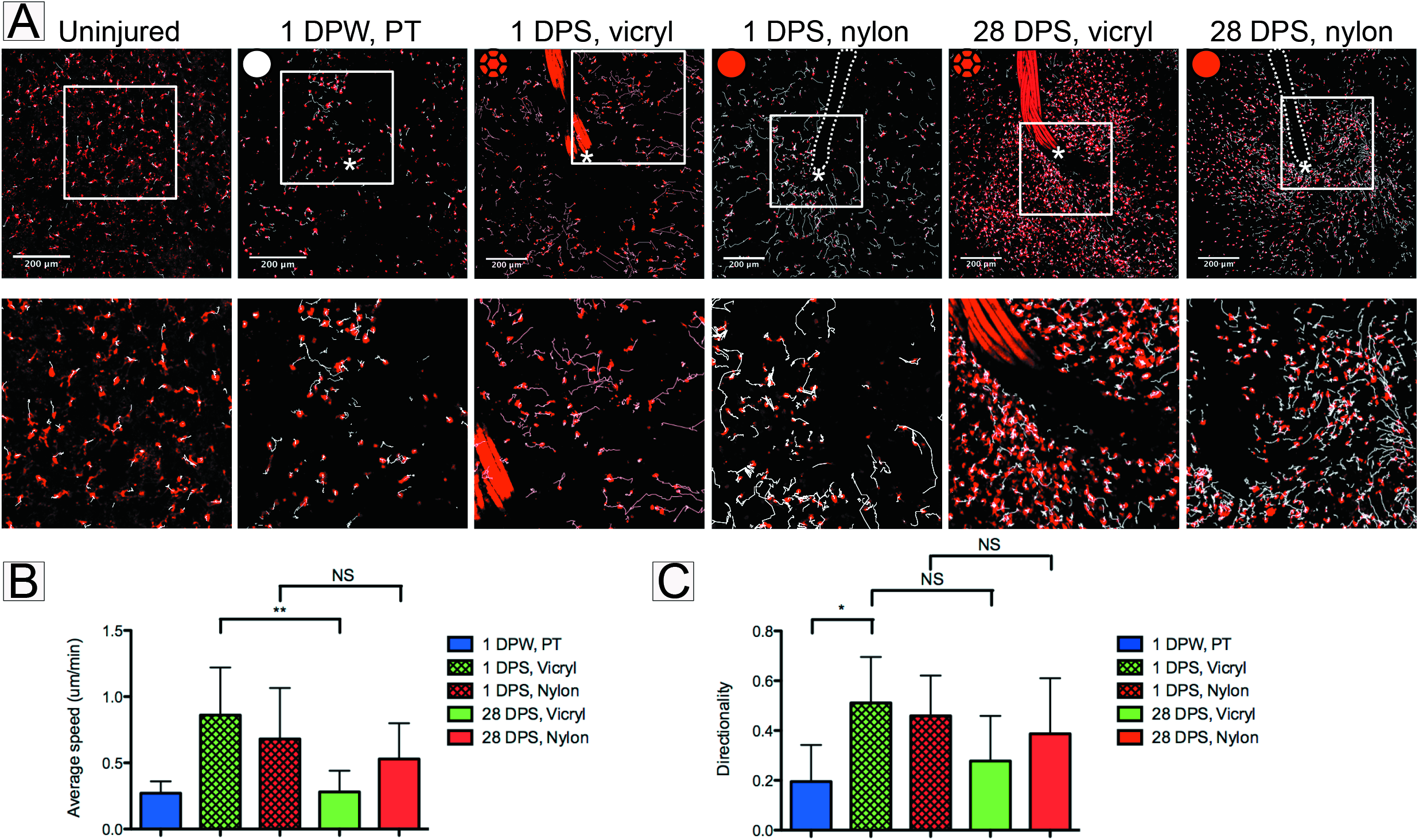
Immune cell motility and directionality within suture-adjacent tissues is affected by implanted material. A) Endpoints from 180 minute long representative timelapse movies of Tg(*mpeg*:mCherry) transgenic adult zebrafish at the indicated timepoints, of uninjured, post pull through or suture implant, showing the tracks of macrophages as they respond to the wound/suture. Tracks are generated by automated cell tracking software (see Methods) and indicated in white; area of wounding or implantation marked by white asterisk; dotted lines indicate nylon suture. N = 4 independent fish per condition, per timepoint. B) Quantification of mean macrophage speed at 1 day and 28 days post implantation, averaged from tracking data, indicating how motility is suppressed at later timepoints, particularly with vicryl sutures. Statistical significance, as measured by one-way ANOVA, is P = 0.0045. C) Quantification of directionality of macrophages at 1 and 28 days post implantation, averaged from tracking data. Statistical significance, as measured by one-way ANOVA, is P = 0.0078. Significance values: *P ≤ 0.05, **P ≤ 0.001. NS = not significant. Data information: error bars indicate mean ± SD. Scale bars: A = 200μm.

### Tissue inflammation is exacerbated by FBR and induces the formation of an avascular region

An effective angiogenic response is pivotal for both wound healing (Eming et al, 2014) and biomaterial integration (Spiller et al, 2015). Our previous work has indicated that pro-inflammatory macrophages expressing tumour necrosis factor α (tnfα) are critical in driving sprouting angiogenesis during tissue repair, but that macrophages must switch to an antiinflammatory, *tnfα* negative state at later stages to enable appropriate subsequent vessel remodelling and regression (Gurevich et al, 2018). The Tg(*tnfα*:GFP) transgenic line has previously been used to identify the pro-inflammatory state of cell lineages other than leukocytes including intestinal epithelial cells during inflammatory bowel disease (Marjoram et al, 2015). We used our suture implantation model to observe the dynamic changes that occur with respect to both tissue inflammation and angiogenesis during a FBR. We combined the Tg(*tnfα*:GFP) transgenic line that marks pro-inflammatory cells with the Tg(*mpeg*:mCherry) macrophage marker line to reveal macrophages with pro-inflammatory or anti-inflammatory phenotypes. Acute (pull through suture) wounds and both suture types showed increased numbers of pro-inflammatory macrophages at early timepoints following insult, with numbers of these cells resolving back to uninjured levels by 28 DPW in acute wounds and nylon sutures (Figure 4A). Acute wounding (pull through sutures) further reveals that the broader wound *tnfα* expression response is also transient, peaking at 7 DPW, being largely resolved by 14 DPW and entirely resolved by 28 DPW (Figure 4A, B), as previously described for acute wounds (Gurevich et al, 2018; Hubner et al, 1996; MacLeod & Mansbridge, 2016). By contrast, both nylon and vicryl sutures maintain a significant level of *tnfα* expression in the vicinity of the foreign body throughout the observed 28 days (Figure 4A, B). The overall *tnfα* response induced by vicryl sutures extends out to a much larger area compared with the nylon suture, indicating that tissue inflammation varies with respect to the nature of the implanted material.

**Figure 4:**
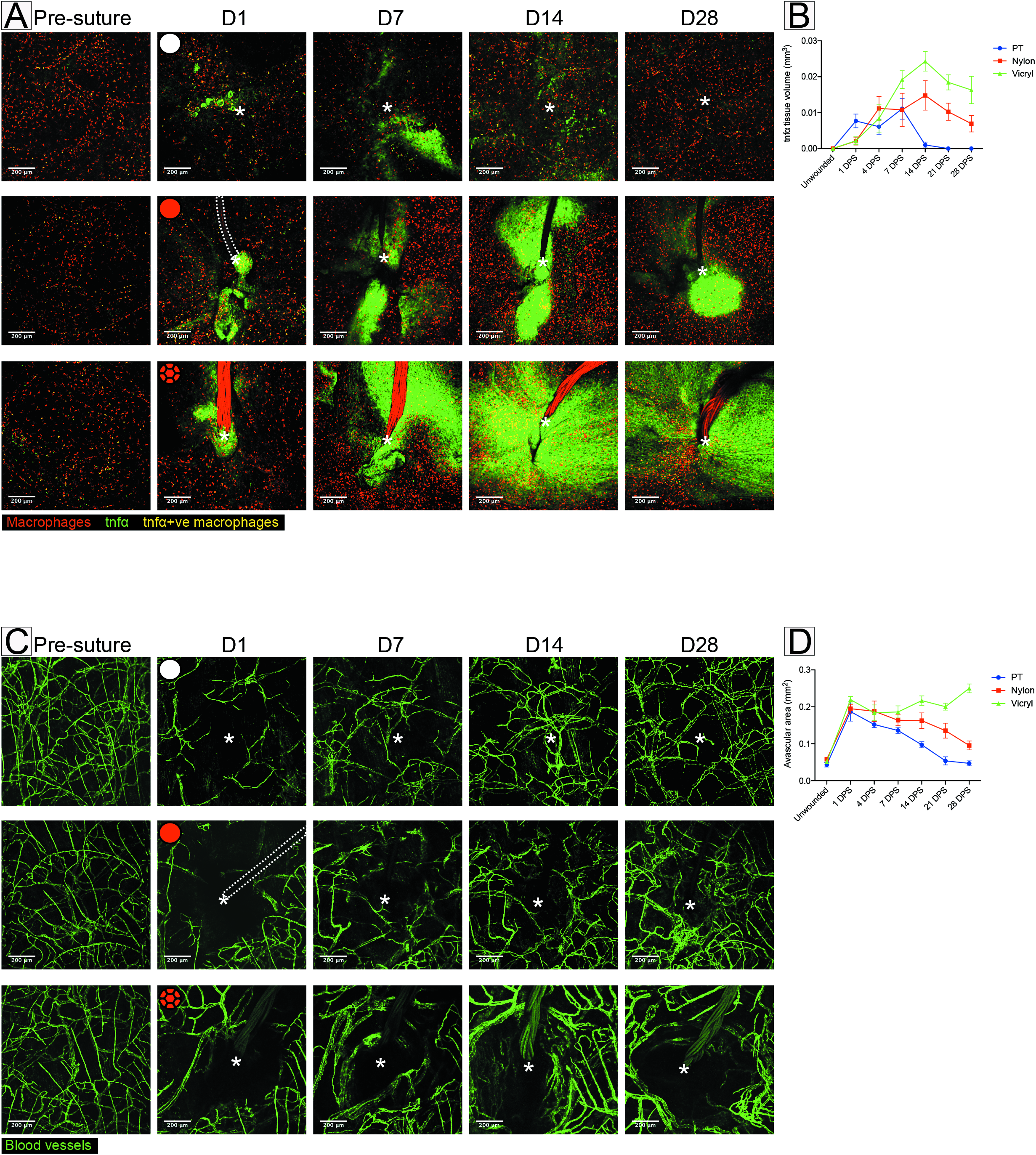
Extent of *tnfα* expression and size of avascular zone are also dependent on suture type. A) Representative images of Tg(*tnfα*:GFP); Tg(*mpeg*:mCherry) double transgenic zebrafish immediately prior to and following suture pull through, or implantation of nylon or vicryl suture, at indicated timepoints, showing macrophages (red), pro-inflammatory macrophages (yellow), and stromal cells expressing *tnfα* in the vicinity of the wound/suture zone (green). Area of wounding or implantation marked by white asterisk; dotted lines indicate nylon suture. N = 8 independent fish per condition. B) Quantification of total inflammatory area surrounding the wound/suture, measured from images in A). C) Representative images of Tg(*fli*:GFP) transgenic zebrafish immediately prior to and following suture pull through, versus implantation of nylon or vicryl suture, to reveal angiogenic response at the indicated timepoints. Area of wounding or implantation marked by white asterisk; dotted lines indicate nylon suture. N = 8 independent fish per condition. D) Quantification of the extent of avascular zone immediately adjacent to wound/suture, measured from images in D). Data information: error bars indicate mean ± SD. Scale bars: A = 200μm, D = 200μm.

To examine the angiogenic response to suture implantation, we utilised the Tg(*fli*:GFP) transgenic line that marks all blood vessels (Lawson & Weinstein, 2002). Acute wounds (pull through sutures) showed a robust revascularisation response, which was well underway by 14 DPW and largely completed by 28 DPW (Figure 4C, D), in line with the progressive reduction of tissue inflammation (Figure 4A, B). By contrast, both suture types displayed a reduced capacity to establish blood vessels within close proximity of the implantation site, leading to an avascular zone forming around the suture (Figure 4C, D). However, the dynamics of this avascular zone differed for the two suture types: nylon sutures appeared to largely re-establish a vascular supply right up to the suture by 28 DPS, whereas vicryl sutures exhibited a progressive increase in the avascular zone, extending out to 350μm from the suture at 28 DPS. Together, these results suggest a close association between inflammation and subsequently impaired angiogenesis, with the extent of both dependent on the type of implanted biomaterial.

### Dampening the inflammatory response results in reduced fibrosis and improved revascularization

Several studies have examined the relationship between extended, ‘chronic’ inflammation in the context of impaired healing and how this leads to progressive fibrosis (Morais et al, 2010). To test whether this correlation might be causal in FBR, we next attempted to modulate the inflammatory response to suture implants. Our first manipulation utilized the *csf1ra* mutant (referred to as *panther*) that has previously been shown to suppress the normal wound inflammatory response (Gurevich et al, 2018) in combination with Tg(*tnfα*:GFP); Tg(*mpeg*:mCherry) transgenic lines. The *panther* mutant led to a significant “rescue” of the chronic inflammatory state, with reduced expression of *tnfα* in the tissues adjacent to both nylon and vicryl sutures, more closely resembling that of acute pull through wounds at all timepoints (Figure 5A, B) than the equivalent tissues in WT fish (compare to Figure 4 A, B). Combining the *panther* mutant with Tg(*fli*:GFP) revealed a rescue of the avascular zone defect also, with vessels now growing considerably closer to the implanted suture (Figure 5C, D; compare to Figure 4 C, D).

**Figure 5:**
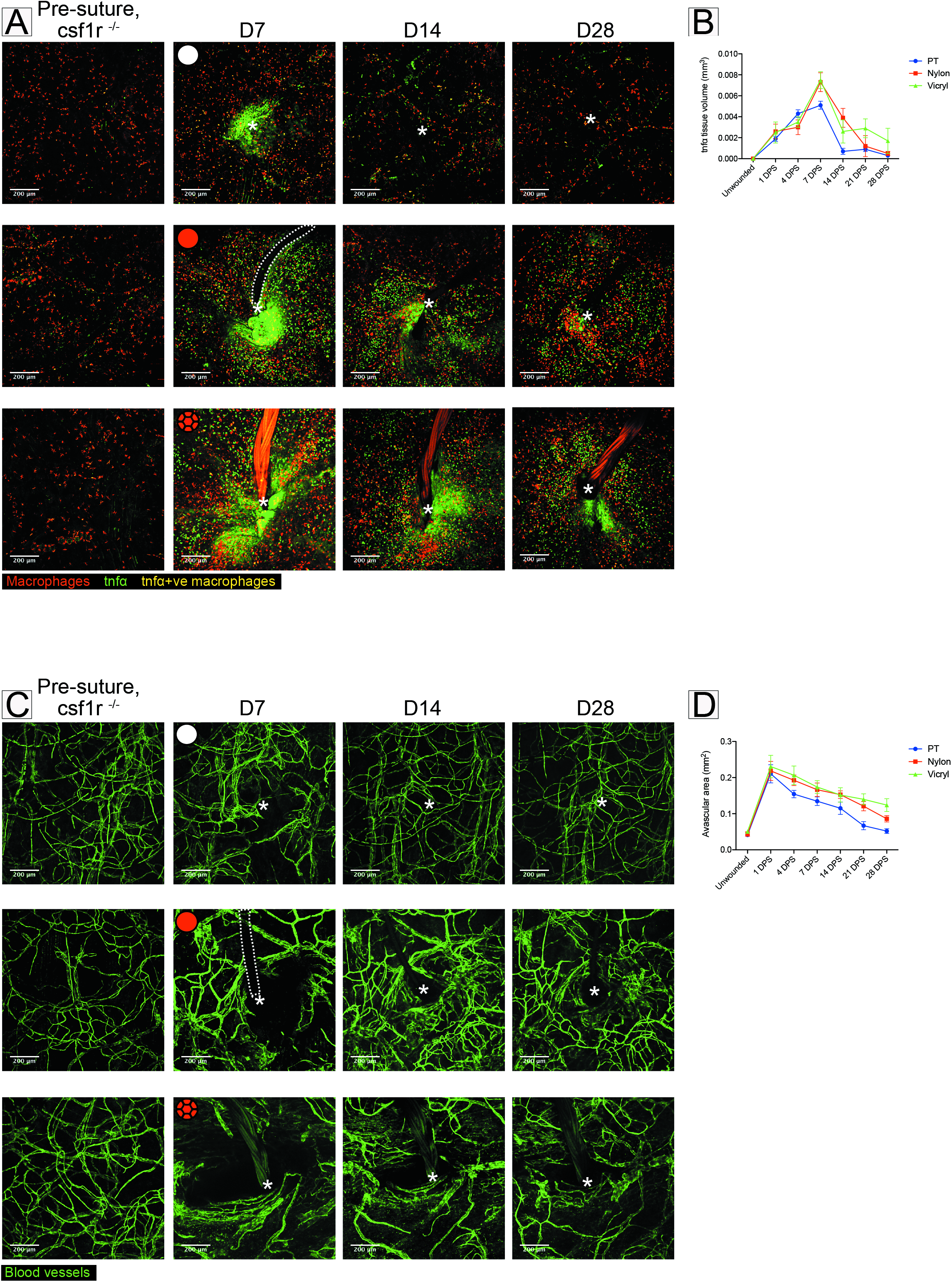
Genetic immunosuppression results in decreased TNFα expression and reduced avascular zone. A) Representative images of *csf1ra* ^−/−^ Tg(*tnfα*:GFP); Tg(*mpeg*:mCherry) double transgenic adult zebrafish immediately prior to and following suture pull through, or implantation of nylon or vicryl suture, at indicated timepoints. Area of wounding or implantation marked by white asterisk; dotted lines indicate nylon suture. N = 6 independent fish per condition. B) Quantification of the altered inflammatory area surrounding the wound/suture, measured from images in A). C) Representative images of Tg(*fli*:GFP) transgenic adult zebrafish immediately prior to and following suture pull through, or implantation of nylon suture or vicryl suture, at indicated timepoints. Area of wounding or implantation marked by white asterisk; dotted lines indicate nylon suture. N = 6 independent fish per condition. D) Quantification of avascular zone immediately adjacent to wound/suture, measured from images in C). Data information: error bars indicate mean ± SD. Scale bars: A = 200μm, E = 200μm.

To complement this genetic approach we also suppressed inflammation in wild type fish by treatment with hydrocortisone (from 7 days to 28 days post suture implantation – Figure 6A), similar to treatment regimes used in previous zebrafish studies (Hasegawa et al, 2017; Richardson et al, 2013). This treatment lead to a comparable rescue of FBR and restoration of tissue repair and biomaterial integration to that seen in the *panther* mutant scenario, with much reduced avascular and fibrotic zones (Figure 6B-E). Furthermore, dampening of inflammation also lead to fewer FBGCs and a decrease in the average volume of observed macrophages (Figure 6F, G; compare to Figure 2 C). Together, these two strategies identify chronic tissue inflammation as a likely candidate for driving the FBR process and, by implication, leading to failure of biomaterial integration, suggesting that the ability to dampen tissue inflammation might be a valuable tool in the amelioration of these problems in a clinical setting.

**Figure 6:**
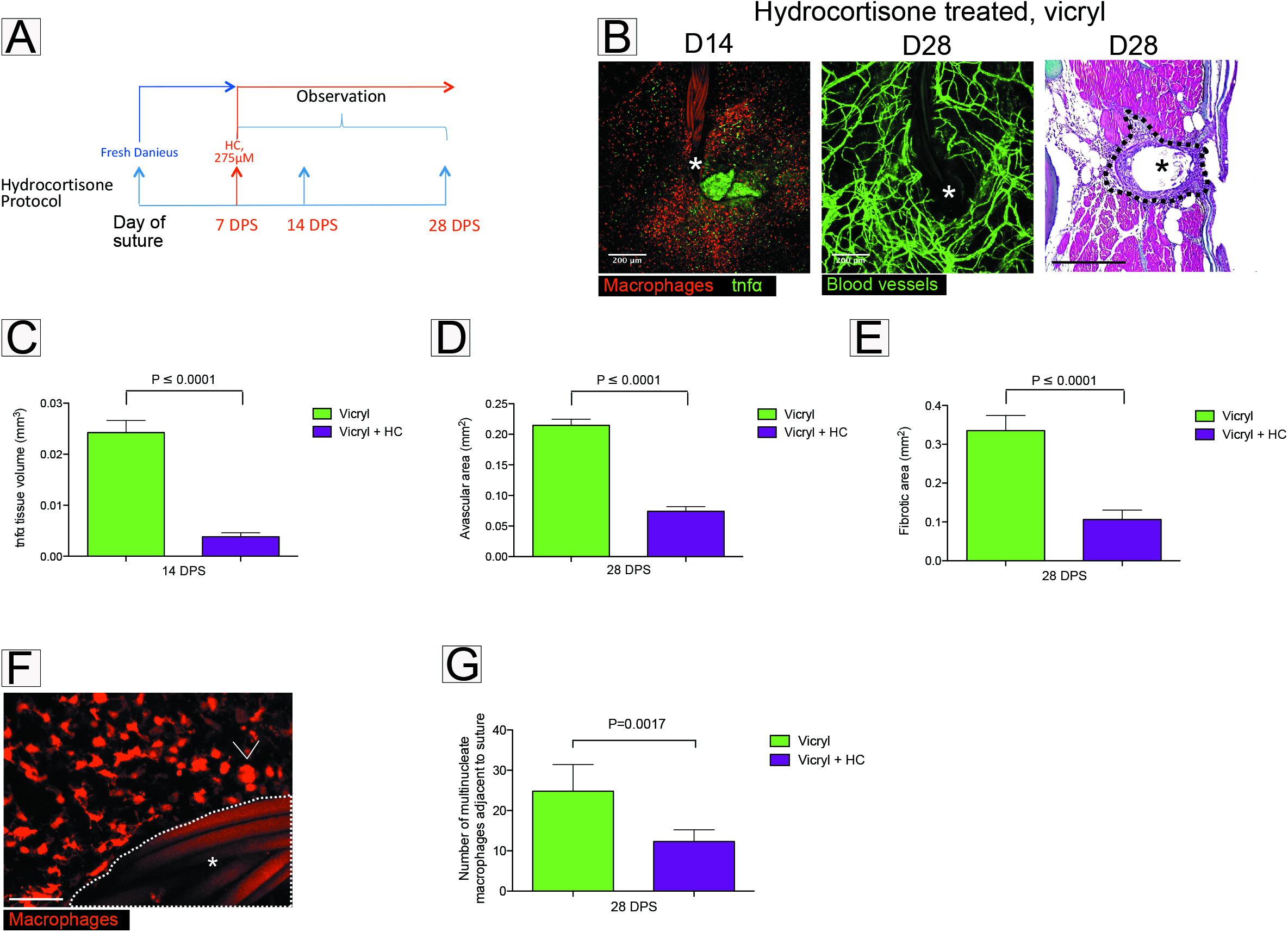
Pharmacological anti-inflammatory intervention results in a reduced avascular zone, fewer foreign body giant cells and decreased fibrotic encapsulation. A) Diagram showing Hydrocortisone treatment protocol used to dampen inflammation during FBR (7-28 DPS). B) Representative images of Tg(*tnfα*:GFP); Tg(*mpeg*:mCherry), Tg(*fli*:GFP) suture site and Masson’s trichrome stained section of suture tissue at the indicated timepoints following vicryl suture implantation and treatment with Hydrocortisone. Site of suture implantation marked by asterisk. N = 6 independent fish per condition. C) Quantification of total inflammatory area surrounding the wound/suture, measured from images as in B). Statistical significance, as measured by two-tailed t-test, is P ≤ 0.0001. D) Quantification of total area of fibrotic encapsulation, measured from images as in B). Statistical significance, as measured by two-tailed t-test, is P ≤ 0.0001. E) Quantification of avascular zone immediately adjacent to wound/suture, measured from images as in B). F) Representative image of Tg(*mpeg*:mCherry) transgenic fish at 28 DPS, showing considerably fewer FBGCs (arrowheads) adjacent to vicryl suture (white asterisk) following Hydrocortisone treatment. N = 5 independent fish per condition. G) Quantification of images from F, showing that average number of FBGCs adjacent to vicryl suture (within 200μm radius) decreases following Hydrocortisone treatment. Statistical significance, as measured by two-tailed t-test, is P = 0.0008. Data information: error bars indicate mean ± SD. Scale bars: B = 200μm, F = 50μm.

## Discussion

Our new model of the foreign body response to suture implantation in zebrafish has allowed us to observe the dynamic interplay of inflammation on cells and tissues including the vasculature and stromal cells that deposit collagen at the implantation site. These studies imply a direct relationship between the extent of the inflammatory response and the degree of fibrotic encapsulation of a foreign body such as a suture and have several implications for the clinic.

### A reciprocal relationship between inflammation, angiogenesis and scarring

Our previous work, as well as that of others, has characterized a dramatic angiogenic response at sites of acute tissue damage. This results in a transient increase in vessel density in the vicinity of the damage site to fuel increased metabolic requirements as the wound heals. These vessels subsequently regress, remodel and normalize back to that seen in uninjured tissue as the repair process finishes (Gurevich et al, 2018; Johnson & Wilgus, 2014). This tightly regulated wound angiogenic response is presumed to be critical because failed angiogenesis associates with chronic, non-healing wounds (Demidova-Rice et al, 2012; Nunan et al, 2014). Interestingly, our current study indicates that biomaterial implantation leads to an avascular zone, which correlates closely with the extent of tissue inflammation, and subsequently also with the zone of fibrotic encapsulation that occurs as a consequence of the FBR. This avascular zone has also been observed in response to implantation of other biomaterials such as biosensors, and is known to impair the integration and function of such devices (Morais et al, 2010). Indeed, revascularization post implantation is considered a key element in determining whether a biomaterial integrates or fails (Morais et al, 2010; Yu et al, 2009).

Avascular zones are not an entirely pathological phenomenon; cartilage is avascular, as is the zone beneath the developing epidermis of embryonic skin. Establishment of these avascular territories does not involve inflammation and is believed to be due to presence of avascular glycosaminoglycans such as Hyaluronic Acid (Feinberg & Beebe, 1983; Hallmann et al, 1987; Martin, 1990). A better understanding of which signals drive the avascular zone in the context of a FBR, and whether they are directly or indirectly released by inflammatory cells, may guide us towards ways for improving vascularity and better tissue integration with implanted biomaterials.

### Modulating the inflammatory cells to regulate angiogenesis and fibrosis

We have previously demonstrated that pro-inflammatory macrophages that form the first wave of an acute inflammatory response following wounding upregulate vascular endothelial growth factor (VEGF) and are important in driving sprouting angiogenesis (Gurevich et al, 2018); it is also clear that these pro-inflammatory cells induce collagen deposition and fibrosis at repairing wound sites (Cash et al, 2014; Cash & Martin, 2016; Wynn & Barron, 2010). Our current study reveals that implantation of a foreign material triggers general tissue inflammation, which varies in extent depending on the type of material. Intriguingly, this zone of inflammation is closely correlated with both the observed fibrotic encapsulation response and the subsequent avascular zone. We have previously observed this potential link between fibrosis and vascularization in the context of osteopontin knockdown, with the suppression of this wound inflammatory marker in mouse wounds resulting in reduced scarring, increased angiogenesis and rapid repair (Mori et al, 2008). In our present work, reduction in tissue inflammation by genetic means (*panther* mutant) or chemically (hydrocortisone treatment) appears to decrease both fibrosis and avascularity, suggesting that inflammation might be a critical factor in determining whether the biomaterial will integrate or undergo rejection via FBR, given the association between implant failure, extent of angiogenesis and fibrosis. Indeed, this is in line with recent attempts to make biomaterials more biocompatible that have focused on reducing inflammation (Kim et al, 2017; Zhang et al, 2014). However, our results also indicate that pro-inflammatory macrophages are still present in areas immediately adjacent to implanted sutures, particularly up to 14 days post suture implantation; this corresponds with the main vessel sprouting response during the repair and integration process, suggesting that these cells are likely playing a similar important pro-angiogenic role as previously determined (Gurevich et al, 2018). Taken together, our results suggest that future innovations to maximize biomaterial integration should distinguish between pro-inflammatory macrophages and other pro-inflammatory tissues.

### What role for giant cells in the FBR?

Foreign body giant cells (FBGCs) which are presumed to be formed by macrophage fusion, were first described 50 years ago (Mariano & Spector, 1974) and are considered a characteristic component of the tissue response to implanted materials as well as to some parasitic infections (Chambers, 1978; Davis et al, 2002). Indeed, activation and aggregation of distinct, specialized macrophages in response to persistent and antagonistic stimuli such as mycobacterium are now thought to be the key events driving the formation of granulomas in response to TB (McClean & Tobin, 2016; Ramakrishnan, 2012). Our study is the first to dynamically image these cells aggregating in high densities prior to FBGC formation. We note that FBGCs are more commonly seen in the vicinity of vicryl versus nylon sutures, suggesting that there may be a threshold level for both cell density and phenotypic state of inflammatory response before fusion will occur. Macrophages that come into direct contact with certain biomaterials are believed to undergo a process of ‘frustrated phagocytosis’, where the inability to engage with the material drives the fusion process; this leads to a subsequent decrease in phagocytic ability and a concomitant increase in free radical, enzyme and acid release to degrade implanted materials (Anderson et al, 2008). However, many questions remain concerning the precise triggers and mechanisms that underlie this fusion process, to activate FBGC formation. Our model presents a valuable opportunity for unraveling the dynamic nature of these fusion mechanisms, and gaining insight into what the specific function of FBGCs may be.

### Adult zebrafish as an important new *in vivo* model of FBR and clinical implications

This study represents the first time that the cell and tissue interactions underlying the FBR between biomaterial and surrounding tissue have been imaged dynamically, *in vivo* and non-invasively. The numerous similarities to mammalian FBR marks the zebrafish as a valuable model for increasing our understanding of the cellular and molecular basis for FBR in response to specific biomaterials. Our approach is particularly powerful as it allows the examination of several key processes and cell players – inflammation, formation of FBGCs, and the angiogenic response – in the same animal over time, permitting the specific tracking and dissection of dynamic cell:cell conversations. In addition, the imaging opportunities in zebrafish combined with its genetic tractability and amenability for chemical/pharmacological intervention, have allowed us to investigate how modulating inflammation in various ways may impact on tissue restoration during FBR. The advances in our understanding presented here will drive further identification and refinement of methods that alleviate FBR and improve biomaterial integration.

**Sup. Figure 1:** Representative Masson’s Trichrome stained transverse sections of pull through, nylon or vicryl sutured fish at 4, 7, 14 and 28 DPS, as indicated. Site of wound or suture indicated with black asterisk. N = 5 independent fish per condition per timepoint. Scale bars = 200μm.

**Sup. Figure 2:** A) Several z-stack images from a single “large” macrophage adjacent to a vicryl suture from the image shown in Figure 2D (boxed area), with arrows indicating multiple nuclei (green) within the cell.

**Movie 1:** Slice by slice z-stack movie of the cells shown in Figure 2D, identifying numerous macrophages as FBGCs with multiple nuclei. Movie z-step size = 2μm, scale bar = 20μm.

**Movie 2:** Representative z-projection timelapse movie of the flank of a laterally mounted uninjured Tg(*mpeg*:mCherry) transgenic, showing tracking of macrophage dynamics and ‘patrolling’ behaviour, imaged every 5 minutes for 180 minutes.

**Movie 3:** Representative z-projection timelapse movie of the flank of a laterally mounted ‘pull through’ injured Tg(*mpeg*:mCherry) transgenic at 1DPW, showing tracking of macrophage dynamics, imaged every 5 minutes for 180 minutes.

**Movie 4:** Representative z-projection timelapse movie of the flank of a laterally mounted, vicryl sutured Tg(*mpeg*:mCherry) transgenic, 1DPS, showing tracking of macrophage dynamics, imaged every 5 minutes for 180 minutes.

**Movie 5:** Representative z-projection timelapse movie of the flank of a laterally mounted, nylon sutured Tg(*mpeg*:mCherry) transgenic, 1DPS, showing tracking of macrophage dynamics, imaged every 5 minutes for 180 minutes.

**Movie 6:** Representative z-projection timelapse movie of the flank of a laterally mounted, vicryl sutured Tg(*mpeg*:mCherry) transgenic, 28DPS, showing tracking of macrophage dynamics, imaged every 5 minutes for 180 minutes.

**Movie 7:** Representative z-projection timelapse movie of the flank of a laterally mounted, nylon sutured Tg(*mpeg*:mCherry) transgenic, 28DPS, showing tracking of macrophage dynamics, imaged every 5 minutes for 180 minutes.

## Materials and Methods

### Zebrafish strains and maintenance

All experiments were conducted with approval from the local ethical review committee at the University of Bristol and in accordance with the UK Home Office regulations (Guidance on the Operation of Animals, Scientific Procedures Act, 1986). Wild type and transgenic lines Tg(*fli1*:eGFP) [referred to as Tg(*fli*:GFP)](Lawson & Weinstein, 2002), Tg(*mpx*:GFP)(Renshaw et al, 2006), Tg(*mpeg1*:mCherry) [referred to as Tg(*mpeg*:mCherry)](Ellett et al, 2011), TgBAC(*tnfα*:GFP) [referred to as Tg(*tnfα*:GFP)](Marjoram et al, 2015) were maintained on TL wild type background, and staging and husbandry were performed as previously described (Westerfield, 1995). The mutant strain used was *csf1ra^j4e1^* (Parichy et al, 2000), maintained on AB background or used in combination with transgenic lines as indicated. *csf1ra^j4e1^* mutants were genotyped by visual inspection for absence of mature xanthophores as previously described (Parichy et al, 2000).

### Suture implantation into adult zebrafish

Adult zebrafish suturing was performed as previously described (Witherel et al, 2017). Briefly, zebrafish were anesthetized in tank system water with 0.1mg/mL tricaine (ethyl 3-aminobenzoate methanesulfonate, Sigma Aldrich, Hamburg, Germany) and subsequently placed onto a foam surgical stand for surgery. Single interrupted sutures were implanted by placing a single loop through the tail, approx. 3mm anterior to the tailfin, using either nylon non-absorbable sutures (Polyamide, 8-0 monofilament, 3/8 tapered needle, S&T, Neuhausen, Switzerland) or vicryl, absorbable sutures (Polyglactin, 8-0 braided, 3/8 needle, Ethicon, Somerville, NJ, USA). Pull through control wounds were generated by implantation of a suture at the exact same anatomical location, which was immediately ‘pulled through’ and removed.

### Imaging of adult zebrafish

For all imaging experiments, fish were initially anaesthetized in 0.3% Danieau’s solution with 0.1mg/mL tricaine, and subsequently imbedded in a 10cm petri dish, using 1.5% w/v agarose added over the tail. Care was taken to keep agarose away from the gills. For timelapse imaging, fish were maintained in a lightly anaesthetized state at 0.05mg/mL tricaine throughout to allow continued breathing; fish that were no longer breathing by the end of movie acquisition were excluded from analysis. Gross anatomical images were generated on a Leica M205 FA system (Leica Microsystems). Confocal images and timelapse movies were generated on a Leica SP8 MP/CLSM system (Leica Microsystems).

### Hydrocortisone treatment

For drug treatments, fish were treated with 275μM Hydrocortisone (Sigma Aldrich) dissolved in ethanol, as previously described (Richardson et al, 2013). 0.1% absolute ethanol was used for all treatments as well as vehicle control.

### Masson’s Trichrome staining

Harvested fish tails were immediately fixed in 4% PFA overnight at 4°C on a rocker, washed with PBS and then decalcified in 0.5M ethylenediaminetetraacetic acid (EDTA) (Sigma Aldrich, Hamburg, Germany) for seven days at 4°C on a rocker, replacing the EDTA solution on the third day. Samples were then stained for Masson’s Trichrome, as previously described (Witherel et al, 2017).

### Transmission Electron Microscopy

Tails were harvested at 28 DPS, fixed and processed as previously described (Nunan et al, 2015). Ultrathin (0.02 μm) sections were images on a Tecnai 12-FEI 120 kV BioTwin Spirit transmission electron microscope.

### Image analysis and statistics

All image analysis was performed in ImageJ. Detection, tracking and spatial analysis of immune cells used the Modular Image Analysis automated workflow plugin for Fiji (Cross, 2019; Rueden et al, 2017; Schindelin et al, 2012). Sample motion due to tissue growth was corrected using translation-based registration via the SIFT Align plugin for Fiji (Lowe, 2004; Saalfeld, 2008) followed by B-spline unwarping using the BUnwarpJ plugin (Arganda-Carreras et al, 2006). Cell features were enhanced prior to detection using the WEKA pixel classification plugin (Arganda-Carreras et al, 2017). Noise in the enhanced image was removed using a 3D median filter and immune cells isolated from background using the Otsu threshold method with a constant user-defined offset (Otsu, 1979). The binarised image was refined using 2D hole filling and a 3D distance-based watershed transform (Legland et al, 2016). Immune cells were identified in 3D as contiguous regions of pixels labelled as foreground using the MorphoLibJ plugin (Legland et al, 2016). An outline of the suture was manually annotated, with any immune cells detected coincident with it assumed to correspond to misdetection; these cells were excluded from further analysis. Immune cells were tracked between frames using the Munkres algorithm with scores based on object centroid separation (Munkres, 1957). Track spatial coordinates were used to calculate instantaneous velocity and track orientation in the XY-plane. A static reference point corresponding to the suture was manually-identified in each video (separate from the previously-detected outline). The angle between the instantaneous track orientation and this point was also measured (i.e. an angle of 0° corresponds to a cell moving directly towards the suture).

All statistical analysis was performed using Graphpad Prism. Data was confirmed to be normally distributed via d’Agostino-Pearson test or Shapiro-Wilk test prior to further comparisons. Student’s t-test were used except in the case of comparisons involving more than two groups; in these instances, one-way ANOVA was performed for all comparisons, and a Bonferroni multiple comparison test was subsequently performed.

