## Supplementary figures and images for "Live imaging the Foreign Body Response reveals how dampening inflammation reduces fibrosis"

### Supplemental Figure 1

#### Sup. Figure 1

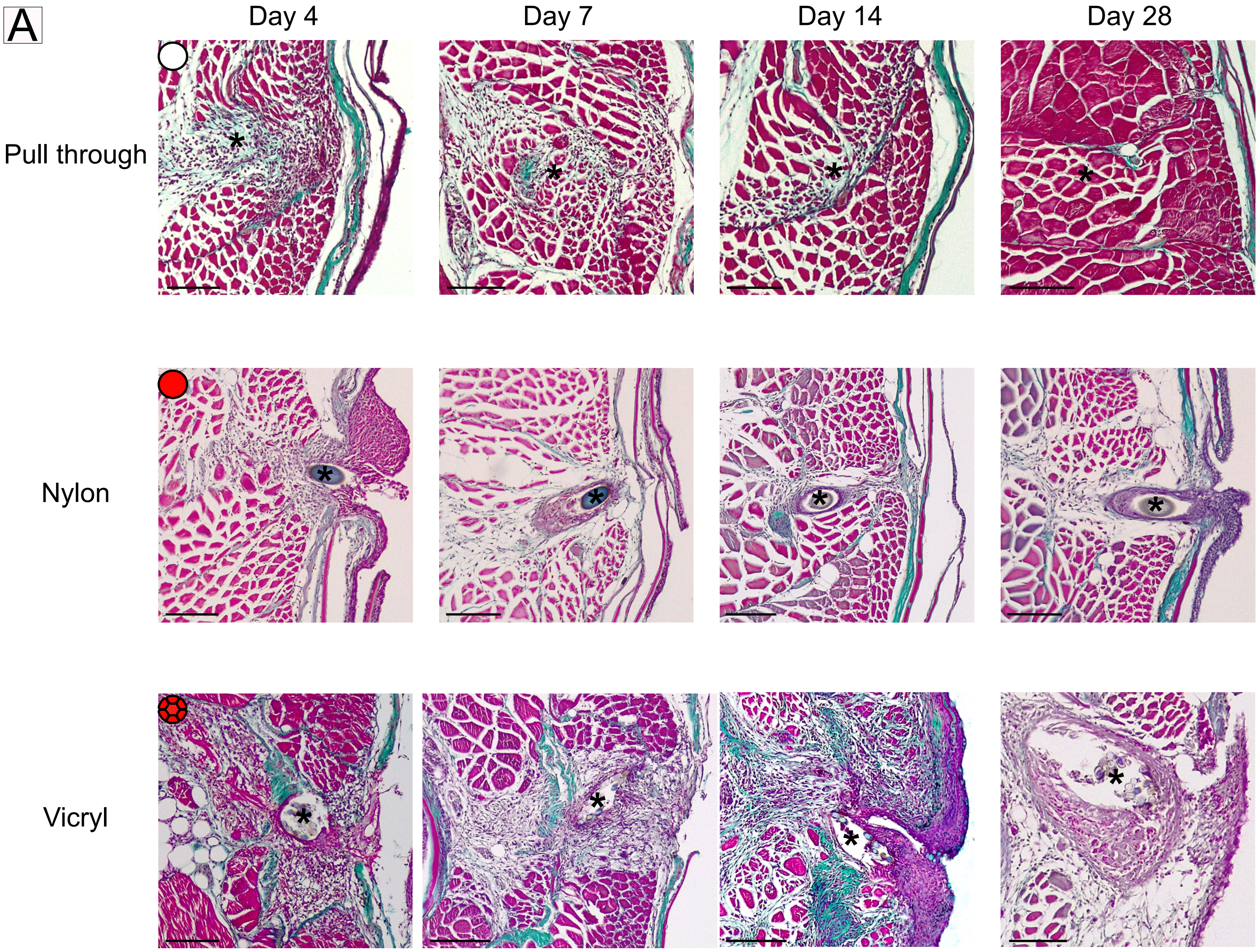

### Supplemental Figure 2

#### Sup. Figure 2

A

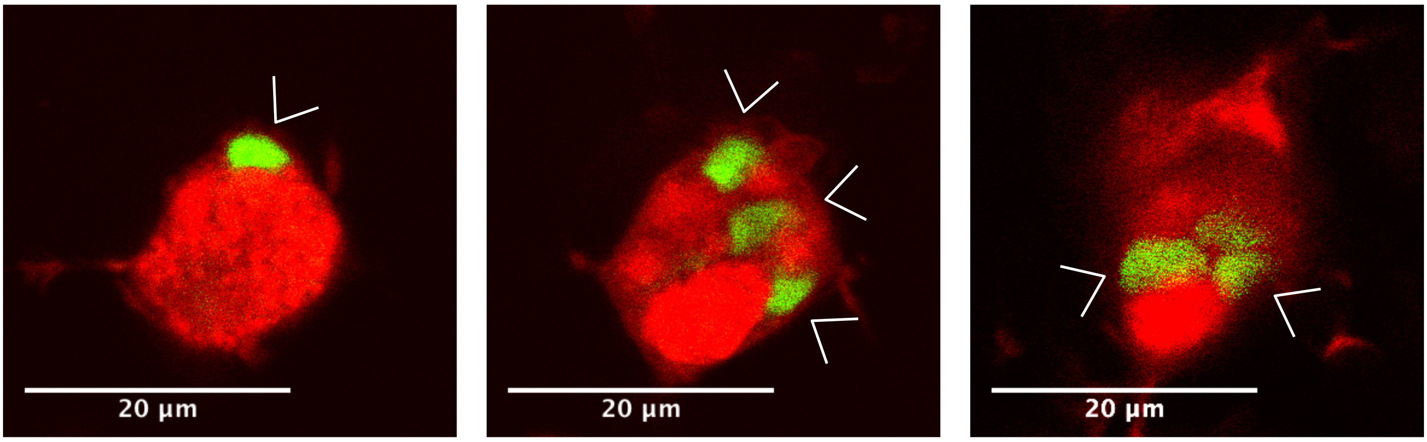
